# No division of labour, and subfertile foundresses, in a phyllode-gluing Acacia thrips

**DOI:** 10.1101/755371

**Authors:** James D. J. Gilbert

**Affiliations:** School of Environmental Sciences, University of Hull, Cottingham Road, Hull HU6 7RX, UK; Fowlers Gap Arid Zone Research Station, School of Biological, Earth & Environmental Sciences, University of New South Wales, NSW 2052, Australia

**Keywords:** Pleometrosis, joint nesting, cooperation, quasisocial, parasocial, division of labour

## Abstract

Behavioural variation is a hallmark of animal societies, which commonly contain breeders and nonbreeders, and helpers and nonhelpers. In some cases labour is divided with nonbreeders “helping” – gaining indirectly, via genetic benefits, or directly, e.g. by augmenting group size. Conversely, they may benefit by not helping, conserving energy for breeding later. However, subordinate behaviour after inheriting a breeding position is rarely evaluated.

In the Australian interior, Acacia thrips *Dunatothrips aneurae* (Thysanoptera) glue *Acacia* phyllodes together into “domiciles”. Foundresses, usually sisters, build domiciles singly or communally. Some co-foundresses are nonreproductive, and their role is currently unknown. I experimentally rejected the idea that they substantially “help” by contributing to domicile repair. Nonreproductives were less likely to repair damage than reproductives. Alternatively, they may be waiting to inherit the domicile, or simply of too poor quality to reproduce or help. To test these alternatives, in the field, I allowed repairer or nonrepairer females to “inherit” a domicile by removing their nestmate(s). Thus isolated, “nonrepairer” females took much longer to repair domiciles than “repairers”, control singletons or pairs. Although ovarian condition was equivalent across groups, after 21 days nonrepairers actually laid fewer eggs compared to other groups.

Thus, labour was not divided: instead reproduction and helping covaried, probably depending on female quality and the outcome of intra-domicile competition. Nonreproductive nonhelpers were not waiting to breed. Their role, and their net effect on colony productivity, remains to be shown. They are likely subfertile, and may make the “best of a bad job” by gaining indirect benefits to the best of their limited ability.

## INTRODUCTION

Social groups are composed of members that often vary in their contribution to necessary tasks. In cooperative or eusocial societies, at any given time individuals typically partially or completely specialize in particular tasks, especially reproduction (“reproductive skew”; Vehrencamp 1979; Vehrencamp 1983a; Vehrencamp 1983b), but also other tasks (Beshers and Fewell 2001), for example, foraging (Johnson 2010), parental care (Browning et al. 2012), nest defence (Gerber et al. 1988), nest homeostasis (Jandt et al. 2009), and others (see Komdeur 2006). Within these societies, costly non-reproductive tasks such as foraging and defence are typically performed by “helpers” who are completely or partially reproductively suppressed, while breeding is performed by one or a few reproductive individuals (Solomon and French 1997; Koenig and Dickinson 2004).

In some species, however, non-reproductive putative “helpers” do not appear to engage in helping (Cant and Field 2001; Korb 2007; Browning et al. 2012). This can sometimes be explained if the helper in question is waiting to inherit a breeding position and would incur reproductive costs later by helping now (Field and Foster 1999; Cant and Field 2001; Leadbeater et al. 2011). Alternatively, a helper may appear “lazy” compared to other helpers if it has less genetic stake in current offspring (e.g. Curry 1988), may function as an insurance workforce in case other helpers are injured (Baglione et al. 2010) or may be helping in subtle ways that may not have yet been measured (Turnbull et al. 2012; Frank et al. 2018). Perhaps more parsimoniously, one simple and long-standing explanation for non-reproductive, non-helping individuals is that they are simply of poor quality, or subfertile - and, as such, may or may not help depending on the extent of their limited ability, making the “best of a bad job” (Eberhard 1975; Craig 1983). Perhaps surprisingly, there is as yet no unequivocal empirical evidence for the “subfertility hypothesis” (Strassmann and Queller 1989), although there are several cases where it has been shown not to apply (Sullivan and Strassmann 1984; Leadbeater et al. 2011). In one example, Field & Foster (1999) found that nonreproductive worker hover wasps given the chance to “inherit” their nest subsequently developed ovaries and began reproducing.

In contrast to cooperative species in which contributions to different tasks are typically sharply divided, some other groups are characterized by less pronounced behavioural and reproductive specialization, e.g. lions (Packer et al. 2001), rodents (Hayes 2000), burying beetles (Trumbo and Wilson 1993) and some sweat bees (although see Abrams and Eickwort 1981; Kukuk 1992). Where breeding is thus shared, individuals may help according to their genetic stake in a brood (Hoogland 1983; Curry 1988), or conserve their helping effort for a time when they are themselves reproducing (Packer et al. 2001).

*Dunatothrips aneurae* Mound (Thysanoptera:Phlaeothripidae), are tiny thrips that live in the arid Australian Outback, where they can be solitary or social (Morris et al. 2002). Females construct “domiciles” by gluing together terminal phyllodes of *Acacia aneura* trees into a completely enclosed nest, within which they produce a single generation of offspring (Gilbert and Simpson 2013). Offspring develop entirely within the domicile, whose integrity is maintained by resident adults. Domiciles are built from a silklike anal secretion (henceforth “silk”) and function to maintain humidity for developing larvae (Gilbert 2014). Domiciles are initiated by solitary females or by groups of 2-5 (rarely up to 20), typically but not always related (Bono and Crespi 2008). The factors promoting nest co-founding are only partially understood, partly because individual roles within groups are unknown. Co-founding carries survival advantages to all females, especially when invaded by kleptoparasites (Bono and Crespi 2008), but also entails reproductive competition for space which is intensified in small domiciles (Gilbert et al. 2018).

There is a low and variable degree of reproductive competition and skew within cofounded *D. aneurae* domiciles. Cofoundresses vary in ovarian development, with a proportion of females nonreproductive, especially in small domiciles (Gilbert et al. 2018). However, whether reproduction co-varies with any form of “helping” is unknown. Females are uniformly pacifist towards intruders (Gilbert et al. 2012; Gilbert and Simpson 2013) suggesting a lack of any defence-based helping behaviour. On the other hand, the production of silk by nonreproductive females may constitute altruistic behaviour (West et al. 2007). Silk is necessary for domicile function (Gilbert 2014) and its production is potentially costly (see Craig et al. 1999 for a review of “real” silk production in animals).

Any trade-off between silk and egg production would create at least the potential for selection for division of labour within domiciles (Gilbert 2014). During fieldwork, I noticed that cohabiting female *D. aneurae* varied in their latency to begin repairing experimental damage to domiciles - variation that may potentially reveal just such a division of labour. If good repairers are bad reproducers, and *vice versa*, this would be good evidence of cooperative division of labour within *D. aneurae* domiciles. In contrast, if good repairers are good reproducers, and *vice versa, D. aneurae* is more likely to be a communal breeder, where subordinate females either (a) help in proportion to their genetic stake in offspring or, similarly, are waiting to inherit the breeding position; or (b) are of too poor quality either to breed or help at the same level as breeders. These two alternatives can be separated by the behaviour of subordinates that are experimentally allowed to inherit the breeding position: if subordinates fail to begin to breed upon inheritance of the nest, alternative (b) is more likely. I asked two questions:

1. Are reproduction and domicile repair divided among foundresses?
2. Do nonreproductive foundresses breed upon inheritance of the domicile?

To address these questions, I performed two experiments. First, in the lab, I identified “repairer” and “nonrepairer” females by their latency to begin repairing experimental damage, and then investigated whether repair tendency was associated with ovary development and egg maturation, assessed via dissection. Second, in the field I conducted a removal experiment to test whether the tendency to repair the nest was associated with reproduction when females were given the chance to breed within the domicile - both potential reproduction (ovarian status) and realized reproduction (eggs laid). I did this by removing all other females, allowing the focal female to “inherit” the domicile.

I had no particular *a priori* expectations with which to form strong predictions. On the one hand, in species that form associations of foundresses from the same generation, which is almost certainly true for *D. aneurae*, strong reproductive skew and division of labour is rarely found (being more typical of matrifilial associations; Reeve and Keller 1995). Supporting this, morphological castes do not appear to occur in *D. aneurae*. On the other hand, well-studied relatives of *D. aneurae* within the Phlaeothripinae are eusocial (*Kladothrips* spp.; Crespi 1992), although, being gall-inducers and with matrifilial societies, their ecology is somewhat different (Crespi et al 2004). Yet even morphologically similar individuals within groups can often show naturally occurring behavioural variation in task performance, which can self-organize into eusocial-like patterns of division of labour (Page 1997; Jeanson et al. 2005). I therefore made a tentative prediction that I would find at least incipient division of labour within *Dunatothrips* domiciles, such that breeding and repairing would be negatively associated.

## METHODS

Fieldwork was conducted between January and October 2013 at a large patch of *A. aneura* at the Bald Hills paddock (S 30° 57′ 39″ E 141° 42′ 18″) near the University of New South Wales Arid Zone Research Station on the Fowlers Gap property, approx. 110km N of Broken Hill, Australia (see Gilbert and Simpson 2013 for details).

### Are reproduction and repair divided?

For lab studies, I used a subset of the *D. aneurae* domiciles collected from the field and dissected as described in Gilbert & Simpson (2013). In 48 nests, during nest dissections I experimentally damaged the nest with watchmaker’s forceps, and assessed their repair behaviour before removing them from the nest and dissecting them to assess their ovarian (i.e. reproductive) condition. Only dealate (reproductively mature) individuals were included in experimental observations, as typically only these individuals contribute to repair (Gilbert and Simpson 2013). I made a small tear in the nest, roughly equivalent to about 20% of the silk area, and folded back the silk. Typically at least one foundress would thereupon begin repairing immediately (within 30 seconds), which I scored as 2 (“Immediate repair”). I continued these initial observations for 1 h. During this time, individuals were brushed lightly on the pronotum with a few grains of a coloured fluorescent micronized paint powder (US Radium Co®), applied using a fine paintbrush, to create a temporary distinctive ID that allowed me to follow individuals within the nest. After 1 h I removed the first immediate repairer from the nest, using a probe made from the hair of a shaving brush, and observed the behaviour of the remaining female(s) over the subsequent 1 h period. One of more of these females typically began repairing upon removal of a nestmate. I scored these individuals as 1 (“Delayed repair”). Over successive 1 h periods I removed one-by-one the immediate/delayed repairers in the sequence that they began repairing, and observed any changes in behaviour in remaining females. Some individuals remained inactive within the nest for the duration of the experiment, even if another female was repairing damage only a few millimetres away, and never began repairing. I scored these as 0 (“Never repaired”).

Removed individuals were dissected, and the extent of development of their ovaries was examined, according to protocols described in Gilbert et al (2018). I recorded the number and volume (π×length×(width/2)^2^) of any developing oocytes, and also of any mature, chorionated eggs that were ready to lay. I also measured female body size (pronotum width). I tested for differences in ovarian development according to repair tendency using Kruskal-Wallis tests.

### Do nonreproductives breed when they inherit the domicile?

Our removal experiment can be summarized briefly as follows: In domiciles with 2 or 3 individuals, I first categorised all females as “repairer” or “nonrepairer” according to their tendency to repair an experimentally damaged nest. Then I experimentally removed a subset of females from each nest (1 or 2 females per nest, so as to leave 1 behind) and dissected them to assess the reproductive status of their ovaries. I then assigned nests randomly to treatment groups in which I replaced either a repairer (+R group), or a nonrepairer (+NR group), thereby allowing (or forcing) this female to “inherit” her domicile. This remaining female I left in the nest for 21 days to assess (1) her efficacy in repairing the damage, and (2) her realized reproduction over this period. After this period, the remaining female was removed and dissected. The two treatment groups, +R and +NR, were compared to control groups consisting of singly built nests (CS group) and nests built by female pairs (CP group).

I identified *Dunatothrips aneurae* domiciles on thin-phyllode variants of *A. aneura* trees in the field containing either 2 or 3 dealate females, and which were at a stage of near-complete construction (i.e. without eggs or larvae). Foundress females were initially counted within intact domiciles non-invasively, using a powerful LED (LED Lenser®) shone through the silk from behind; these preliminary counts and the absence of eggs were confirmed following experimental nest damage (see below). Where possible, nests were chosen that were simple in shape without convolutions that might complicate experimental removals, or which might prevent individuals from detecting experimental damage to the domicile silk.

To test individual contributions to repair, I slightly damaged the domicile wall by peeling away approximately 10-20% of its surface area using watchmaker’s forceps. I then observed the behaviour of the inhabitants for the next 3 h or until both (or all) individuals had responded by repairing the domicile. Repair behaviour is very obvious in this tiny insect, even in the field, as the female prepares to lay down a strand of silk by vigorously waving her abdomen; the function of this waving is probably either mechanical (pumping the abdomen) or sensory (locating with abdominal hairs the surfaces to be tied together). During this 3 h period, near-constant observation (at least every 30 seconds) allowed me to keep track of individuals without marking them. At the low humidities typical of the Outback environment, 6 h is long enough for small larvae to die of desiccation when exposed outside a domicile (Gilbert 2014), so 3 h represents an amount of time within which repairing a nest can have meaningful effects upon larval survival.

Reflecting my lab classification of repair behaviour, I classified individuals into repairers (R) and nonrepairers (NR). An individual was classified as a repairer if, during the observation period, it added more than a negligible amount of new silk to the damaged area (I disregarded a minority of individuals that added one or a few silk strands and then became quiescent). A two-female domicile could thus be classified as (R,R), (NR, R) or (NR, NR). In practice (NR, NR) domiciles were rare; I only encountered 2 during my fieldwork.

I randomly allocated the domiciles to three experimental groups. In all cases, I initially removed both (or all) individuals from the domicile, and then replaced different individuals depending on the experimental group. Both removal and replacement were done using a coarse hair probe and replacement was completed within 30 minutes of removal. In the “Leave Repairer” group (+R, n=11) I removed both (or all) individuals from the domicile and then replaced one repairer. In the “Leave Nonrepairer” group (+NR, n=8) I instead replaced one nonrepairer. In the control pairs group (CP, n=9) I replaced both (or all) individuals. In the control singletons group (CS, n=46), I identified early-stage colonies that were singly founded and performed the same experimental damage procedure, before removing and replacing the foundress female. All permanently removed individuals were taken at this point to the lab for dissection to determine ovarian development (see above).

Experimental domiciles were monitored each day in the field for 21 days. I noted the time taken to repair the experimental damage by visually categorising the state of repair as “no repair”, “incomplete repair” or “complete repair”. “No repair” was where a negligible number of silk strands had been added to the damaged area and the area remained substantially open to the environment. “Incomplete repair” was judged to be where repair had begun but strands were sparse and the damaged area was either (i) not completely covered by silk strands, or (ii) covered very loosely by silk such that any perturbation of the phyllodes (e.g. by mild wind) would likely reopen the hole in the silk. “Complete repair” was where the new silk completely covered the damaged area to the extent that gentle perturbation of the phyllodes would be unlikely to reopen the hole in the domicile wall. I also recorded any instances of naturally occurring domicile damage (recording the repair of this damage in the same way as above), domicile destruction, or abandonment during the experimental period.

After a period of 21 days, experimental domiciles were collected and taken to the lab. 21 days is the maximum time between domicile completion and egg-laying seen previously in the lab (Gilbert and Simpson 2013), and I assumed an individual’s reproduction in the field during that time would be a reliable, standardized snapshot of its egg-laying rate, as a proxy for fitness. The number of eggs and larvae in each domicile was counted and the latest stage of larvae attained was noted. I excluded from analysis domiciles that were blown down, predated or destroyed over the experimental period.

To analyse both repair times and numbers of offspring, I used generalised linear models (GLM) with poisson errors and “treatment” as a single predictor variable. I compared the treatment groups using planned orthogonal contrasts:

1. Control pairs (CP group) against all singletons (pooled mean of CS, +R and +NR groups).
2. Experimental nonrepairers (+NR group) against all other singletons (pooled mean of CS and +R groups)
3. Experimental repairers (+R group) against control singletons (CS group).

I dissected all removed individuals as described above, recording number and volume of developing oocytes. All individuals remaining in domiciles after the 21 d experimental period were also dissected. Owing to time constraints, only a subset of the CS group (n=8) was used for this part of the experiment.

I asked whether R and NR individuals from experimental domiciles differed in reproductive status, both before (i.e. individuals removed immediately) and after the experiment (individuals left in the domicile for 21 days).

I used linear models of the total volume of developing oocytes (log-transformed after adding the minimum nonzero value in the dataset to all values; the precise value of this small added increment did not affect the results). As predictor variables, I included main effects of repair tendency (R versus NR) and treatment (before versus after), plus their interaction.

The treatment term in this analysis was perfectly confounded by the number of females cohabiting in the domicile at the time of removal (“before” individuals all had >1 females in each domicile at the time of removal, while “after” individuals uniformly had only 1 individual per domicile). My experiment was unable to separate these two confounded variables, but I was able to shed partial light on the problem by conducting a second analysis that also included individuals in the CS and CP groups. All of these individuals had been “left” for the entire experimental period, but they differed in the number of females in the nest: 1 for the CS group and >1 for the CP group. For this second analysis, instead of treatment (“removed” versus “left”), I instead used the number of females in the domicile at the time of removal (“1” versus “>1”) as a predictor variable.

## RESULTS

### Are reproduction and repair divided?

24 females in the lab experiment (14 repairers and 10 nonrepairers) had no developing oocytes and were assigned an oocyte volume of zero. Among the remaining 118 females dissected, the volume of developing oocytes varied from 6.0×10^−4^ to 3.0×10^−2^ mm^3^. 65 females had no mature eggs; among the remainder, the volume of chorionated oocytes varied from 1.6×10^−3^ to 2.8×10^−2^ mm^3^. Pronotum width varied from 240 to 360 μm; body length varied from 1625 to 2422 μm. As in Gilbert et al (2018), body size was not related to reproductive state (Kruskal-Wallis test, H = 0.64, df = 1, p-value = 0.72).

I found that females were more likely to repair (whether immediate or delayed) if they had developing eggs than if they did not, and were much more likely to repair if they had mature, chorionated eggs (X-squared = 9.52, df = 4, p-value = 0.049, Figure 1). Body size was not related to repair tendency (Kruskal-Wallis test, H = 0.40, df = 1, p-value = 0.53).

**Figure 1.**
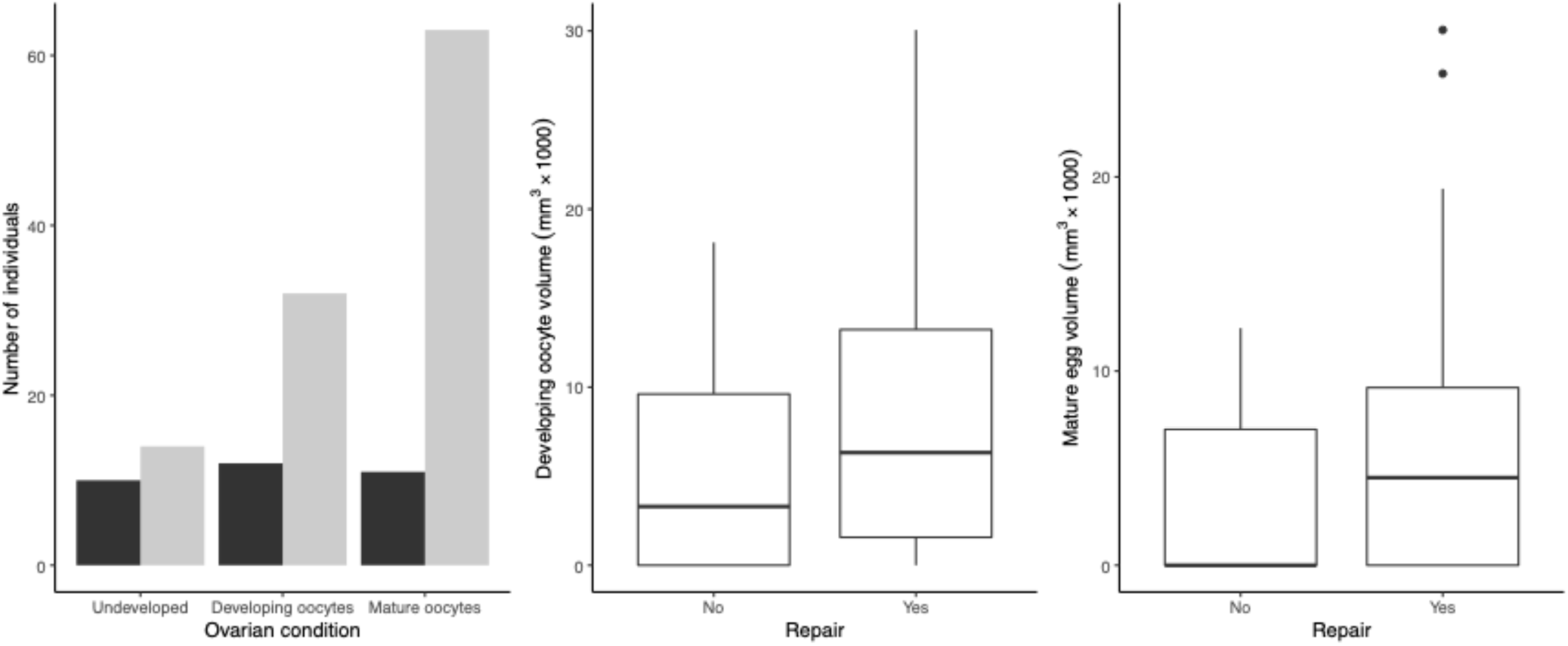
(a) frequency of participation in domicile repair (black = no repair, grey = repair) by female *D. aneurae* in different reproductive condition; (b) Volume of developing oocytes in females according to participation in repair; (c) Volume of mature eggs in females according to participation in repair.

Females that participated in nest repair had only a marginally (nonsignificantly) higher volume of developing oocytes than those that never participated in repair (Kruskal-Wallis test, H=3.41, df=1, p=0.06), and the number of developing oocytes was not associated with repair (Kruskal-Wallis test, H=1.55, df=1, p=0.21). In contrast, females that participated in nest repair had both a higher number (Kruskal-Wallis test, H = 5.24, df = 1, p-value = 0.022) and volume of mature eggs (Kruskal-Wallis test, H = 4.285, df = 1, p-value = 0.038) than those that did not. This pattern was mainly driven by the frequency of females with zero mature eggs: when I restricted the dataset to only those females carrying mature eggs, there was then no difference between repairers and nonrepairers in either number of mature eggs (Kruskal-Wallis test, H= 0.036, df = 1, p-value = 0.85) or volume (H= 0.17466, df = 1, p-value = 0.676).

### Do nonreproductives breed when they inherit the domicile?

Of 21 experimental domiciles with 2 females, 8 were categorised as (R, R), 11 were (NR, R) and 2 were (NR, NR). Of 9 domiciles with 3 females, 1 was (R, R, R), 3 were (R, R, NR) and 5 were (R, NR, NR). All 8 singleton nests in the CS group whose repair activity I assessed in the field were (R). Among repairers, latency to repair varied from <30 seconds to 126 m within the 3 h observation period, while “nonrepairers” were those that failed to repair, or repaired negligibly, within this period.

### Time to repair damage

There were significant differences in repair time among groups (GLM, poisson errors, dropping “group”, Δdeviance=43.76, Δdf=3, p=0.004). These differences were driven entirely by the +NR group (Figure 2a). Domiciles in the control (CP and CS) and +R groups were completely repaired after a median of 2 d following damage, with some completed by the next day. In contrast, the +NR group took a median of 4 d to complete repair (range 2 – 7 d). Contrast 1 was marginally significant (control pairs against all singleton groups: CP against pooled mean of CS, +R and +NR; z=1.651, p=0.099). Contrast 2 was not significant (Experimental repairers, +R group, against control singletons, CS group; z=1.022, p=0.307). Contrast 3 was significant (Experimental nonrepairers, +R group, against pooled mean of CS and +R groups; z=−3.048, p=0.002).

**Figure 2.**
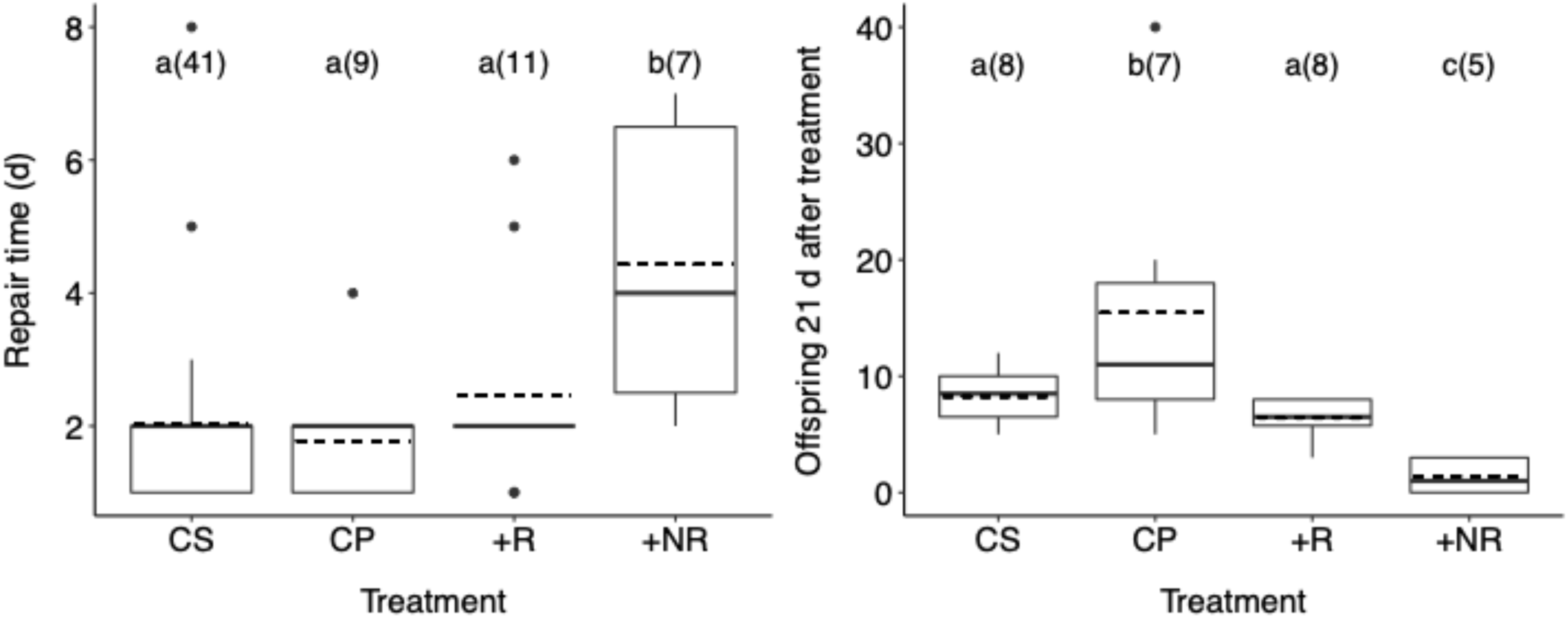
(a) Time to repair damage in *D. aneurae* domiciles from different treatment groups. (b) Number of eggs in the domicile 21 d after treatment. Bars with different letters are statistically different. Sample sizes (domiciles) are given in parentheses. Boxes show median ± IQR, whiskers show range, dashed line shows mean.

### Frequency of domicile damage

There were no differences in the frequency of domiciles destroyed or abandoned during the experimental period: 16 out of 46 domiciles in the CS group, 3 out of 9 domiciles in the CP group, 3 out of 11 domiciles in the +R group and 4 out of 9 in the +NR group (*χ*^2^=0.625, p=0.884).

### Number of offspring

There were significant differences among groups in the numbers of offspring present after 21 days (GLM with poisson errors, dropping “group”, Δdeviance=49.09, Δdf=3, p<0.0001; Figure 2b). CP had more offspring than all other groups (Contrast 1, CP against pooled mean of CS, +R and +NR; z=5.360, p<0.0001). Experimental repairers were similar to control singletons in productivity(Contrast 2, +R against CS; z=−1.383, p=0.167). The +NR group had substantially fewer offspring than all other groups (Contrast 3, +NR group against pooled mean of CS and +R groups; z=−4.225, p<0.0001).

### Reproductive status

Out of the dissected females in my field experiment, 4 females (3 repairers and 1 nonrepairer) had no developing oocytes and were assigned an oocyte volume of zero. Among the remainder (n=64), the volume of developing oocytes varied from 5.3×10^−4^ to 3.4×10^−2^ mm^3^. 22 had no mature eggs, and among the remainder (n=41; 5 females belonging to control groups were not assessed for mature eggs) varied from 5.3×10^−4^ to 3.4×10^−2^ mm^3^.

Individuals of different repair tendency (R versus NR) did not significantly differ in ovarian status, either in the first analysis (using “treatment” as a co-predictor) or in the second analysis (incorporating the CS and CP groups and using “number of females in domicile” as a co-predictor). In the first analysis, there was no interaction between repair tendency and treatment (dropping this interaction, F_1,29_=0.01, p=0.929), nor any statistical effect of repair tendency (dropping repair tendency, F_1,30_=0.179, p=0.676). However, treatment had a significant effect (dropping treatment, F_1,30_=5.99, p=0.020). In the second analysis, again there was no interaction between repair tendency and female number (dropping interaction, F_1,47_=0.217, p=0.643), nor any effect of repair tendency (dropping repair tendency, F_1,48_=0.01, p=0.923). As for “treatment” in the first analysis, dropping female number had a significant upon model performance (dropping female number, F_1,48_=6.55, p=0.014). Individuals sharing domiciles with other females (or which were removed at the beginning of the experiment) had a smaller volume of developing oocytes than females alone in a domicile (or which were removed at the end of the experiment) (Figure 3a). There were no significant differences in volume of mature eggs, whether among treatments or repair tendencies (linear models, all NS; Figure 3b).

**Figure 3.**
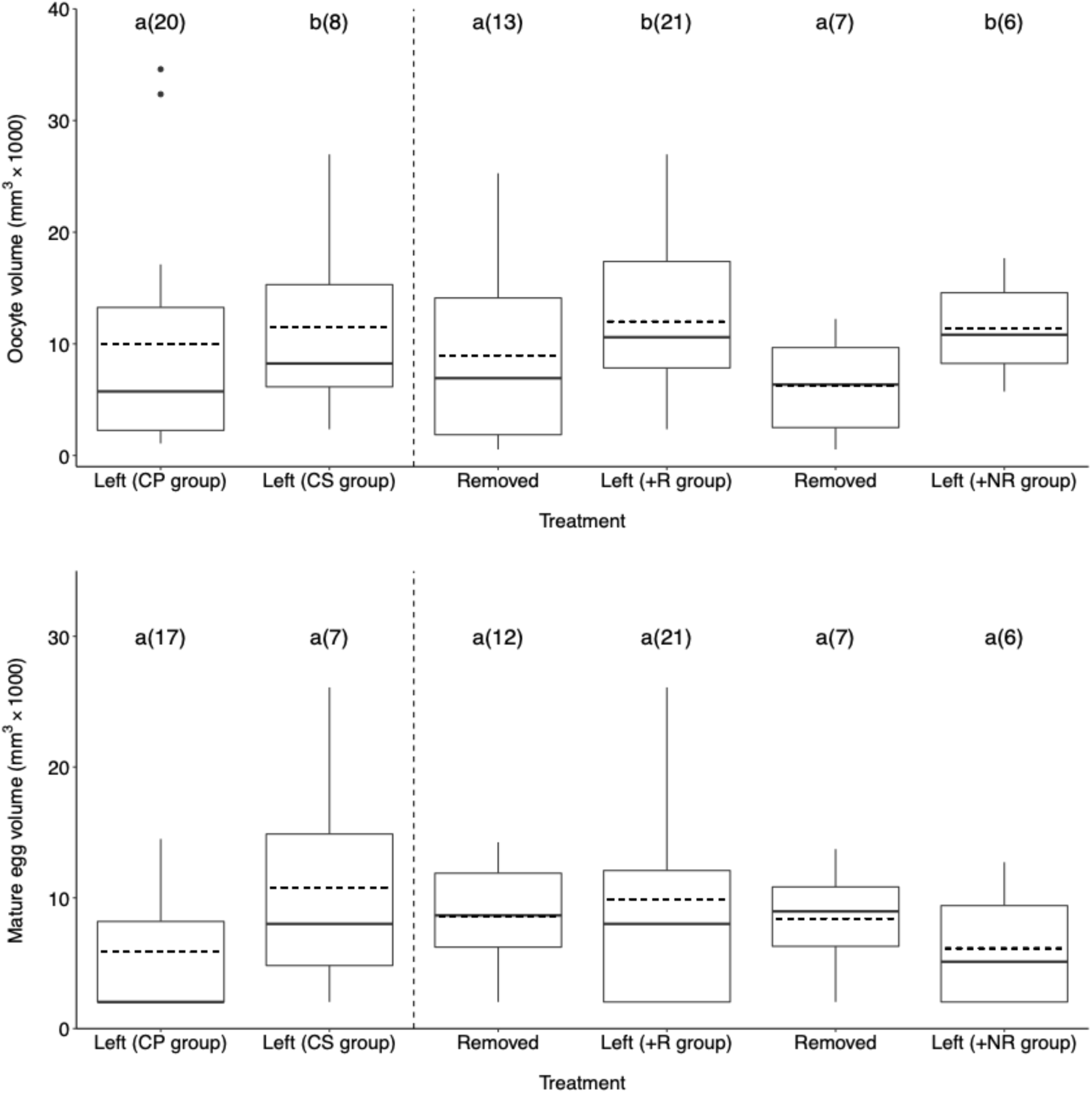
(a) Developing oocyte volume and (b) mature egg volume in females from different treatment groups. Bars with different letters are statistically different. Sample sizes (females) are given in parentheses. Boxes show median ± IQR, whiskers show 95% CI, dashed line shows mean.

## DISCUSSION

In multi-female domiciles of *D. aneurae*, helping effort and realized reproductive output were positively associated: helpers were reproducers, and non-helpers were non-reproducers. Although repairers and non-repairers had statistically similar amounts of developing oocytes both in the lab or in the field, in the lab repairers had more mature eggs, and in the field they ended up actually laying more eggs. A negative association between reproduction and helping would have indicated reproductive division of labour, characteristic of a typically eusocial colony (e.g. Vespid wasps; O’Donnell 1996) or of cooperative breeding vertebrates (Clutton-Brock et al. 2004; Russell et al. 2008).

Instead, with a positive association between realized reproduction and helping, joint-founded *D. aneurae* domiciles resembled low-skew cooperative or communal societies in which group members may help in proportion to their (i) genetic stake in current offspring (Curry 1988; Packer et al. 2001) or probability of future breeding (Hartley et al. 1995; Gilchrist and Russell 2007; Leadbeater et al. 2011), or (ii) quality or condition (as proposed by West-Eberhard 1978; Craig 1983). Between these alternatives, I found little evidence that nonreproductive females were conserving energy for later breeding, suggesting instead that female quality may mediate both helping and breeding effort in this system. Although repairer and nonrepairer females had similar numbers/sizes of developing oocytes (Figure 1b), any given costly task (e.g. domicile repair) may be more costly for females of lower quality, requiring them to resorb oocytes, and consequently they may actually mature and lay fewer eggs (Figure 1c).

Reproductive competition is likely to exacerbate differences among females in realized reproductive output. Competition occurs among *D. aneurae* females, especially in smaller domiciles containing more females (Gilbert et al. 2018) and those which are occupied by inquilines (Gilbert et al. 2012). In smaller domiciles, competition (for example from ecological constraints preventing dispersal, or from successive joining of the domicile by additional females) may force females of poorer quality into a nonreproductive or less reproductive state. For these females, remaining in (or joining) an established domicile may be “the best of a bad job”, so that any small contribution stands a chance of being translated into offspring.

However, breeders gained no apparent advantage by sharing a domicile with nonrepairers. Consistent with Gilbert et al (2018), reproductive females in this study had *fewer* developing oocytes in cofounded domiciles than in singleton domiciles (Figure 3a), suggesting that competition over resources within the domicile reduces fertility of all individuals, rather than nonreproductives being of benefit to breeders, as occurs in some cooperative breeders (Nelson-Flower Martha J. et al. 2013). Furthermore, control pairs and singletons repaired damage equally quickly (Figure 2a). All experimental groups suffered similar rates of wind damage, suggesting that domiciles do not differ in their vulnerability to strong wind, whether maintained by one or two females, or by good versus bad repairers.

It therefore remains unclear whether subfertile females actually help, or whether they reduce group fitness via competition. Research should now focus on testing between these two alternatives. The subfertility hypothesis holds that naturally poor quality females can be selected to invest exclusively in helping, even if their help is less efficient than that of breeders (Eberhard 1975; Craig 1983). Traditionally, the critical test of this hypothesis has been whether putative helpers breed upon experimentally inheriting their natal nest or breeding resource. To my knowledge, there has been no study in which subordinates failed to breed when given this opportunity (e.g. (Field and Foster 1999; Smith et al. 2009; Rehan et al. 2014). If true, this study would provide the first such case. However, previous studies have all been in taxa where subordinates unequivocally act as helpers.

The question of why nonreproductive, apparently costly females should be tolerated by reproductive females, posed and discussed initially by Gilbert et al (2018), therefore remains open. FIrst, it is possible subfertile females may contribute to group fitness in subtle ways not measured here. For example, Bono and Crespi (2006) found that joint-nesting females enjoyed a survival advantage when invaded by kleptoparasitic thrips, providing a possible defensive role for nonreproductive females. Nonreproductives may also participate more actively at the building stage of the domicile than in ongoing maintenance, or may contribute to maintenance of middens in mature domiciles (Gilbert and Simpson 2013). In other social insects, subfertile individuals can have subtle but important effects upon colony function; for example, “soldiers” have been recently discovered to play a primary role in combatting pathogens in both thrips (Turnbull et al. 2012) and ants (Frank et al. 2018). Subordinates may also act via “load lightening”, extending longevity of breeders, even though immediate effects are not obvious (Russell et al. 2007), or via “assured fitness returns” (Gadagkar 1990; Lucas and Field 2011) whereby the effect of a given unit of help in a well-established group, however small, if positive, is worth more to a subordinate’s inclusive fitness than it would have been had she been breeding on her own and therefore likely to fail.

A second alternative is that nonreproductive, nonrepairing female *D. aneurae* may be reproductively suppressed, either behaviourally through aggression (e.g. Kolmer and Heinze 2000) or via pheromonal control or signalling, as in molerats and many social insects (e.g. Keller and Nonacs 1993; Bennett et al. 1996). In some cooperative societies, putative “helpers” actually perform less work than reproductives (Korb 2007; Browning et al. 2012), consistent with the pattern I show here. This hypothesis is also consistent with the fact that nonreproductives tend only to occur as a result of reproductive competition in small domiciles, as shown in Gilbert et al (2018). However, the data shown here suggest this is unlikely. Theoretically, reproductively suppressed individuals should withhold help only to conserve energy if they may inherit a breeding position later (part of the “future fitness” hypothesis; Cant and Field 2001). However, in *D. aneurae*, nonrepairing individuals neither bred nor repaired the domicile after removal of their nestmate (Figure 2). Moreover, aggressive reproductive suppression in *D. aneurae* is unlikely as I have never observed aggression apart from ejecting males after mating (Gilbert and Simpson 2013), although chemical suppression is still possible.

A third and untested possibility remains that nonrepairing females were in ‘akinesis’ or temporary diapause (Varadarasan and Ananthakrishnan 1982), i.e. waiting to reproduce *much* later. This might be the case, for example, if females take a very long time to accumulate resources for producing their relatively very large eggs (suggested by Gilbert et al 2018). If not collected for dissection, nonrepairers may have “come online” at a point more than 21 days after manipulation, and subsequently either dispersed and bred independently or bred within their existing domicile. I regard these possibilities as unlikely. Regarding dispersal, reproductively mature female *D. aneurae* are dealate (i.e. having lost their wings) (Gilbert and Simpson 2013), reducing their capabilities of independent dispersal - all females involved in this study were dealate. Regarding breeding within the domicile, it is very rare to observe females breeding among the remains of an old brood (JDJG, pers. obs.); there is also as yet no evidence for multiple generations of offspring within *D. aneurae* domiciles (see Gilbert and Simpson 2013) so it is equally unlikely that nonreproductives were offspring of foundresses waiting to breed.

Finally, theoretically, public good games models suggest that “laggards” can coexist peacefully (without punishment) with producers in situations where selection acts at the level of the group as well as the individual (Boza and Számadó 2010), an idea with potential in the *Dunatothrips* system where the likelihood of whole-group mortality is likely to be high relative to individual mortality.

Primitively social or communal societies such as I have demonstrated here in *D. aneurae* have sometimes been thought of as an intermediate step in the evolution of eusociality (Lin and Michener 1972) although it has turned out that the two strategies are often phylogenetically distinct (Kukuk 1992; Danforth and Eickwort 1997; but see Richards et al. 2003). Phylogenetic separation appears also to be the rule in Acacia thrips, since the phyllode-tying habit occurs in a clade separate from all other eusocial thrips (Crespi et al. 2004), but to confirm this idea, we would need to investigate the relationship between breeding and helping in relatives of *D. aneurae*, which are almost entirely unstudied. Domicile size and social tendencies vary widely within the genus, with some being obligately solitary (D. gloius, Bono 2007), and others moderately social such as *D. armatus* (Gilbert, unpublished data). The species that most urgently requires study is *D. vestitor*, which is reported to form much larger societies with up to 70 individuals within domiciles that are loosely merged into “supercolonies” (Crespi et al. 2004 and JDJG, pers. obs.).

## Supporting information

Dataset for fig 1 - analysis of repair versus ovarian status

Dataset for fig 2a - analysis of repair time versus treatment

Dataset for fig 2b - analysis of offspring produced versus treatment

Dataset for fig 3a - analysis of developing oocytes versus removal treatment

Dataset for fig 3b - analysis of mature oocytes versus removal treatment

## ACKNOWLEDGEMENTS

I thank Prof. S. J. Simpson for mentorship, guidance and use of his lab and facilities, Dr K Leggett and G. and V. Dowling for helpful discussions, help during fieldwork and use of facilities, Dr LA Mound and Dr A Wells for assistance with fieldwork and helpful discussions, Drs T and M Flower, Dr LE Browning and Dr A Russell for helpful discussions and Dr P. Bierzychudek for supplying micronized paint.

## Notes

#### Summary of Updates

Added supporting datasets

